# Lactate and glycerol-3-phosphate metabolism cooperatively regulate growth and redox balance during *Drosophila melanogaster* larval development

**DOI:** 10.1101/517532

**Authors:** Hongde Li, Kasun Buddika, Maria C. Sterrett, Cole R. Julick, Rose C. Pletcher, Chelsea J. Gosney, Anna K. Burton, Jonathan A. Karty, Kristi L. Montooth, Nicholas S. Sokol, Jason M. Tennessen

## Abstract

The dramatic growth that occurs during *Drosophila* larval development requires rapid conversion of nutrients into biomass. Many larval tissues respond to these biosynthetic demands by increasing carbohydrate metabolism and lactate dehydrogenase (dLDH) activity. The resulting metabolic program is ideally suited to synthesize macromolecules and mimics the manner by which cancer cells rely on aerobic glycolysis. To explore the potential role of *Drosophila* dLDH in promoting biosynthesis, we examined how *dLdh* mutations influence larval development. Our studies unexpectantly found that *dLdh* mutants grow at a normal rate, indicating that dLDH is dispensable for larval biomass production. However, subsequent metabolomic analyses suggested that *dLdh* mutants compensate for the inability to produce lactate by generating excess glycerol-3-phosphate (G3P), the production of which also influences larval redox balance. Consistent with this possibility, larvae lacking both dLDH and G3P dehydrogenase (GPDH1) exhibit developmental delays, synthetic lethality, and aberrant carbohydrate metabolism. Considering that human cells also generate G3P upon Lactate Dehydrogenase A (LDHA) inhibition, our findings hint at a conserved mechanism in which the coordinate regulation of lactate and G3P synthesis imparts metabolic robustness upon growing animal tissues.

## INTRODUCTION

Nearly a century ago, Otto Warburg observed that tumors exhibit high levels of glucose consumption coupled to oxygen-independent lactate production (Warburg, 1956). This metabolic program, which is commonly referred to as the Warburg effect or aerobic glycolysis, has become a focal point of cancer metabolism research (Vander Heiden et al., 2009). The manner by which tumors consume glucose and generate lactate, however, is not unique to either cancer cells or diseased tissue. In fact, the hallmark characteristics of aerobic glycolysis are activated under a variety of normal developmental conditions, such as during maturation of human T cells (Cooper et al., 1963; Pearce et al., 2013; Wang et al., 1976), formation of vertebrate somites (Bulusu et al., 2017; Oginuma et al., 2017), development of muscle tissue (Tixier et al., 2013), activation of hair follicle stem cells (Flores et al., 2017), and *Drosophila* larval growth (Tennessen et al., 2011). Moreover, studies of the mitochondrial pyruvate carrier (MPC1) reveal that forcibly shifting intestinal stem cells towards a more glycolytic state induces over-proliferation in both mice and flies (Bricker et al., 2012; Schell et al., 2017). Overall, these examples illustrate how the coordinate regulation of glycolytic flux and lactate metabolism play central roles in biomass production, cell fate decisions, and developmental growth (Miyazawa and Aulehla, 2018).

While the exact reason for why cells activate aerobic glycolysis *in vivo* remains debatable, one likely explanation revolves around the redox challenges imposed upon highly glycolytic cells (Vander Heiden et al., 2009). Under conditions of elevated glucose catabolism, glyceraldehyde-3-phosphate dehydrogenase (GAPDH) transfers electrons to NAD^+^, resulting in the formation of NADH. These reducing equivalents must be efficiently removed from NADH because the resulting decrease in NAD^+^ availability can dampen glycolytic flux and restricts growth. LDH relieves this redox burden by coupling NADH oxidation to lactate formation, thus ensuring that NAD^+^ is regenerated at an adequate rate. Therefore, highly glycolytic cells, whether in diseased or normal tissues, become reliant on LDH to maintain redox balance. This hypothesis has long been attractive to the cancer metabolism field because LDH inhibitors could hypothetically interfere with tumor growth while having lesser impacts on normal tissues (Avi-Dor and Mager, 1956). As a result, much of our understanding regarding how LDH influences biosynthesis, growth, and cell proliferation is derived from cancer cells studies.

The goal of using LDH inhibitors to disrupt tumor growth has a rich history rooted in the observation that the pyruvate analogs, such as oxamate, inhibit the growth of HeLa cells in glucose-rich media (Goldberg and Colowick, 1965; Goldberg et al., 1965). More recent analyses support these early studies, demonstrating that both RNAi knockdown of *LDHA* transcripts and LDHA inhibitors disrupt cell proliferation in culture and interfere with tumor growth in mouse xenograft experiments (Billiard et al., 2013; Boudreau et al., 2016; Daniele et al., 2015; Fantin et al., 2006; Qing et al., 2010). Moreover, a conditional *Ldha* mutation prevents the formation of *KRAS-* and EGFR-induced non-small cell lung cancer in mice, thereby providing *in vivo* evidence that some tumors require LDHA (Xie et al., 2014).

Despite the ability of LDHA inhibitors to disrupt the growth and tumorigenicity of certain cancer cells, a growing body of evidence suggests that animal cells can compensate for the loss of LDHA activity. Pancreatic cancer cell lines can become resistant to the LDHA inhibitor GNE-140 by increasing oxidative phosphorylation (OXPHOS) (Boudreau et al., 2016). Similarly, human colon adenocarcinoma and murine melanoma cell lines that lack both LDHA and LDHB increase OXPHOS and are capable of forming tumors in xenograft experiments (Zdralevic et al., 2018). However, the most significant evidence that cellular metabolism readily adapts to the loss of LDH activity is not based on cancer studies, but instead stems from a rare inborn error of metabolism known as glycogen storage disease Type XI (GSD-XI), which results from loss-of-function mutations in the human *LDHA* gene (Maekawa et al., 1990). Other than reports of skin lesions and symptoms associated with exercise intolerance (Kanno et al., 1980; Yoshikuni et al., 1986), GSD-XI patients develop and grow normally (Kanno et al., 1988) which is surprising given the role of LDHA in several developmental and physiological processes. The mild symptoms experienced by GSD-XI patients not only raise the possibility that LDH inhibitors might be ineffective in a clinical setting, but also suggest that studies of animal development can identify the metabolic mechanisms that function redundantly with LDH. Toward this goal, we examined the metabolic consequences of mutating *dLdh* (FBgn0001258) in the fruit fly *Drosophila melanogaster*.

Similar to cancer cells, *Drosophila* larvae increase glycolytic metabolism and dLDH activity as a means of supporting the ~200-fold increase in body mass that occurs during this developmental stage (Graveley et al., 2011; Li et al., 2013; Rechsteiner, 1970; Tennessen et al., 2011; Tennessen et al., 2014b). To determine if this increase in dLDH activity is necessary to maintain redox balance and promote biomass accumulation, we examined how *dLdh* mutations influence larval development. We found that although *dLdh* mutants exhibit a decreased NAD^+^/NADH ratio, this metabolic insult had no noticeable effect on either growth rate or biomass accumulation. Instead, metabolomic analysis revealed that *dLdh* mutants up-regulate G3P production, which also promotes NAD^+^ regeneration and potentially supports larval growth despite loss of dLDH activity. We observed a similar result in *Gpdh1* (FBgn0001128) mutants, which develop normally despite a decreased NAD^+^/NADH ratio. Larvae that lack both dLDH and GPDH1, however, exhibit severe growth defects, developmental delays, and synthetic lethality, thus demonstrating that these two enzymes cooperatively support larval growth. Considering that both cancer cells lines and humans GSD-XI patients also increase G3P production in response to the loss of LDH-A (Billiard et al., 2013; Boudreau et al., 2016; Miyajima et al., 1995), our findings suggest that fundamental aspects of this metabolic relationship are similar in both flies and humans.

## MATERIALS AND METHODS

### *Drosophila* husbandry and genetic analysis

Fly stocks were maintained at 25°C on Bloomington *Drosophila* Stock Center (BDSC) food. Larvae were raised and collected as previously described (Li and Tennessen, 2017). Briefly, 50 adult virgin females and 25 males were placed into a mating bottle and embryos were collected for four hours on a 35 mm molasses agar plate with a smear of yeast on the surface. Collection plates were stored inside of an empty 60 mm plastic plate and placed in a 25°C incubator for 60 hours. All mutations and transgenes were studied in a *w^1118^* background. *Gpdh1* and *dLdh* mutations were maintained in trans to the balancer chromosomes *CyO, p{GAL4-twi.G}, p{UAS-2xEGFP}* (BDSC Stock 6662) and *TM3, p{Dfd-GMR-nvYFP}, Sb^1^* (BDSC Stock 23231), respectively. Unless noted, *dLdh* mutant larvae harbored a trans-heterozygous combination of *Ldh^16^* and *Ldh^17^ (Ldh^16/17^*) and a *dLdh* precise excision control strain (*dLdh^prec^*) were used in all experiments (for a description of these alleles, see Li et al., 2017). *dLdh* mutant phenotypes were rescued using the previously describe transgene {*pdLdh*} (Li et al., 2017). RNAi experiments were conducted using transgenes that target either *dLdh* (BDSC stock 33640) or GFP (BDSC stock 41556).

### Generation of *Gpdh1* mutations

*Gpdh1* mutations were generated using a CRISPR/Cas9 approach (Gratz et al., 2013; Sebo et al., 2014). Two oligos encoding guide RNA sequences that targeted either exon 3 (5’-GGCTTCGACAAGGCCGAGGG −3’) or exon 4 (5’-GATCTGATCACGACGTGTTA −3’) were inserted into the BbsI site of pU6-BbSI-gRNA (Addgene). Each gRNA construct was independently injected into BDSC Stock 52669 (*y^1^ M{vas-Cas9.S}ZH-2A w^1118^*) by Rainbow Transgenic Flies (Camarillo, CA). The mutations *Gpdh^A10^* (19 bp deletion within exon 3; 5’-TCGACAAGGCCGAGGGCGG-3’) and *GpdhB^18^* (7 bp deletion with exon 4; 5’-ACGTGTT-3’) were isolated using a PCR-based sequencing approach. All experiments described herein used a trans-heterozygous combination of these two alleles (*Gpdh*^A10/B18^).

### Generation of the *UAS-Gpdh1* transgene

The *UAS-Gpdh1* transgenic strain was generated by PCR amplifying the *Gpdh1* cDNA from *Drosophila* Genomics Resource Center (DGRC) cDNA clone FI03663 using the oligos 5’-AGAATTCATGGCGGATAAAGTAAAT-3’ and 5’-AGCGGCCGCTTAAAGTTTTGGCGACGG-3’. The *Gpdh1* PCR product was inserted into the EcoRI and NotI sites of pUAST-attB (DGRC) and the resulting plasmid was injected into BDSC Stock 24867 (*M{vas-int.Dm}ZH-2A, PBac{y[+]-attP-3B}VK00031*) by Rainbow Transgenic Flies (Camarillo, CA).

### Gas Chromatography-Mass Spectrometry (GC-MS) analysis

*dLdh* mutants and precise excision controls were analyzed using four independent targeted GC-MS-based metabolomic analyses. Samples were collected, processed and analyzed at either the University of Utah metabolomics core facility or the Indiana University Mass Spectrometry Facility as previously described (Cox et al., 2017; Li and Tennessen, 2018). Each sample contained 25 mid-L2 larvae. For all experiments, six biological replicates were analyzed per genotype. GC-MS data was normalized based on sample mass and in internal succinic-d4 acid standard. Each experiment was statistically analyzed using Metaboanalyst (metaboanalyst.ca) version 4.0 with Pareto scaling (Chong et al., 2018).

### Liquid Chromatography-Mass Spectrometry (LC-MS/MS) analysis

For both *dLdh* mutants and precise excision controls, 100 mid-L2 larvae were collected in a 1.5 mL microfuge tube. Each sample was immediately washed three times using ice-cold PBS, all wash solution was removed, and the sample tube was drop frozen in liquid nitrogen. Metabolite extraction and LC-MS analysis was performed by the University of Utah Metabolomics core facility as previously described (Bricker et al., 2012; Cox et al., 2017). Data was analyzed using a two-tailed student t-test.

### Colorimetric metabolite assays

Glycogen, triglyceride (TAG), trehalose, and soluble protein were measured in mid-L2 larvae using previously described methods (Tennessen et al., 2014a). Briefly, 25 mid-L2 larvae were collected from the surface of a molasses egg-laying cap that contained ~1.5 grams of yeast paste and placed in a 1.5 mL microfuge tube. For each assay, at least six biological replicates were collected from independent mating bottles. Samples were washed three times using ice-cold phosphate buffered saline (PBS; pH 7.4). All PBS was removed from the samples and larvae were homogenized in the appropriate assay buffer. 10 μL of larval homogenate was removed for measuring soluble protein using a Bradford assay and the remaining homogenate was immediately heat-treated at 70°C for 5 minutes. Heat-treated samples were frozen at −80°C until analyzed using the appropriate assay.

NAD^+^ and NADH were measured using the Amplite fluorimetric NAD^+^/NADH ratio assay kit (AAT Bioquest, Inc; 15263) according to instructions. Ten mid-second instar (L2) larvae were washed with cold PBS and homogenized with 100 μL of the lysis buffer. The lysates were centrifuged for 10 min at 3000 *g*, and the supernatants were collected for the analysis. Fluorescence was monitored with a Cytation 3 plate reader (BioTek) at Ex/Em=540/590 nm. The concentrations of NAD^+^ and NADH were normalized to the soluble protein concentrations.

All metabolite measurements were repeated a minimum of three times with six independent samples analyzed per genotype. Data was analyzed using a two-tailed student t-test.

### ^13^C-based metabolic turnover rate analysis

For measurement of metabolic turnover rates from glucose to lactate and glycerol-3-phosphate, mid-L2 larvae were fed with Semi-defined Medium (Backhaus et al., 1984) containing 25% D-glucose-^13^C_6_ for 2 hours, then metabolites were detected using an Agilent 6890N Gas Chromatograph with a 5973 Inert Mass Selective Detector and Gerstel MPS2 autosampler. The isotopologue distributions were corrected based on the natural abundance of elements. The metabolic turnover rate *f_x_* was estimated based on the formula *X^L^/X^T^* = *p(1−exp(-f_x_*t/X^T^)*, where *X^L^* is the amount of ^13^C labeled metabolite (m+3), *X^T^* is the amount of total metabolite pool, *p* is the percentage of glucose-^13^C_6_.

### Graphical Representation of Metabolite Data

All figures were generated using Graphpad Prism 7.0 (version 7.0c). Metabolomic data are presented as box plots, with the box extending from the 25^th^ to 75^th^ percentile, the line in the middle representing the median value.

### Quantification of Mitochondrial Genome Copy Number

Quantitative PCR was used to measure relative ratio of mitochondrial DNA/genomic DNA in precise excision controls and *dLdh* mutants based on a previously described strategy (Oliveira and Kaguni, 2011). Total DNA was isolated from 25 mid-L2 larvae using the Qiagen Core Gene 2 extraction kit. For mitochondrial DNA measurements, DNA samples were diluted 1:100 and the *Drosophila* mitochondrial gene *mt:CoI* was amplified using the oligos 5’-TGCTCCTGATATAGCATTCCCACGA-3’ and 5’-TCCACCATGAGCAATTCCAGCGG-3’. The relative abundance of genomic DNA in the samples was measured by amplifying the *Rpl 32* genomic locus using oligos 5’-AGGCCCAAGATCGTGAAGAA-3’ and 5’-TGTGCACCAGGAACTTCTTGAA-3’.

### Larval Respiration Studies

We quantified routine metabolic rate in precise excision controls and *dLdh* mutants as CO_2_ production using established flow-through respirometry protocols for larval *D. melanogaster* (Hoekstra and Montooth, 2013; Hoekstra et al., 2013). We measured metabolic rate for twenty biological replicates per genotype. Each biological replicate consisted of ten mid-L2 larvae that were placed in a small cap containing 0.5 mL of fly food inside of a glass respirometry chamber. The amount of CO_2_ produced by the group of larvae was measured by flowing CO_2_-free air through the chambers at a rate of 100 ml/min and measuring the CO_2_ produced as a result of metabolism using an infrared CO_2_ analyzer (Li-Cor 7000 CO_2_/H_2_O Analyzer; LI-COR, Lincoln, NE). Each run of the respirometer used a multiplexed system (Sable Systems International, Henderson, NV) to cycle through four chambers that contained larvae and a fifth baseline chamber that were all housed in a thermal cabinet maintained at 26°C (Tritech™ Research, Los Angeles, CA). Genotypes were randomly assigned to chambers within each run. Within each run, two technical replicate measurements were performed for each group of larvae. Technical replicate measures were strongly correlated (*r* = 0.935). We calculated VCO_2_ as the average rate of CO_2_ produced across the 10 min. time interval for the first replicate measure in each run for each biological sample after correcting for any minimal drift in the baseline signal. Each group of larvae was massed using a Cubis^®^ microbalance (Sartorius AG, Göttingen, Germany) before being placed in the respirometer. This allowed us to statistically account for the relationship between mass and metabolic rate when testing for differences between genotypes using a Type II Model regression implemented with smatR (Warton et al., 2006) in the R statistical package (Team, 2017). There was no significant difference between genotypes in the slope relating log(mass) and log(metabolic rate) (i.e., the mass-scaling exponent) (*P* = 0.099); we then tested whether genotypes differed in metabolic rate across masses (i.e., for a difference in the y intercept of the relationship between mass and metabolic rate).

### Larval central nervous system (CNS) and gastrointestinal (GI) tract staining

CNS and intestine dissection and analysis were performed as previously described (Luhur et al., 2017; Luhur et al., 2014). Briefly, size- and age-synchronized larval CNSs and larval intestines were dissected in ice cold 1XPBS and fixed at room temperature for 45 minutes in 4% paraformaldehyde (Electron Microscopy Services) in 1X PBS with and without 0.3% Triton X-100, respectively. For Dpn staining, CNSs were subsequently washed with 0.3% PBT and blocked (1XPBS, 0.5% BSA) for at least 1 hour at room temperature. Primary antibody staining was performed with guinea pig anti-Dpn (provided by James Skeath, 1:500) for 5 hours at room temperature. Secondary antibody staining was performed overnight with AlexaFlour-488 conjugated goat anti-guinea pig antibodies (Life Technologies, 1:1000) and DAPI (0.5 μg/ml) at 4°C. Larval intestines were washed with 0.1% PBT and incubated with DAPI for 15 minutes. Following these secondary washes, CNSs and intestines were mounted in Vectashield mounting medium (Vector Laboratories).

For EdU staining, dissected larval CNSs were immediately incubated in Grace’s insect media supplemented with 1mM EdU. Subsequent manipulations were performed as described in the product manual (ThermoFisher C10338).

For quantifications, multiple Z-steps of individual brain lobes or posterior midguts were acquired using the Leica SP5 confocal microscope in the Light Microscopy Imaging Center at Indiana University, Bloomington. The maximum projection of each Z-stack was generated using FIJI. Dpn and EdU positive cell numbers were manually counted using Adobe Photoshop. The area of larval enterocyte nuclei was measured using FIJI.

## RESULTS

### dLDH maintains larval NAD^+^ redox balance

To understand how lactate synthesis influences *Drosophila* larval metabolism, we used LC-MS/MS to measure the NAD^+^/NADH ratio in larvae harboring a *trans-heterozygous* combination of the previously described *dLdh* loss of function alleles, *dLdh^16^* and *dLdh^17^*, as well as a precise-excision control strain, *dLdh^prec^* (Li et al., 2017). Consistent with a model in which dLDH regulates larval redox balance, our analysis revealed that *dLdh^16/17^* mutant larvae exhibit a decreased NAD^+^/NADH ratio (Figure 1A; Supplemental Table 1). In contrast, the ratios of NADP+/NADPH, reduced glutathione/oxidized glutathione (GSSG/GSH), and ADP/ATP were similar in both mutant and control larvae (Figure 1A; Supplemental Table 1). The abundance of AMP relative to ATP, however, was slightly elevated in *dLdh* mutants (Figure 1B; Supplemental Table 1). Overall, our results demonstrate that loss of dLDH activity interferes with the NAD^+^/NADH balance of larvae raised under standard culture conditions but has minimal effects on other aspects of redox metabolism and energy production.

**Figure 1.**
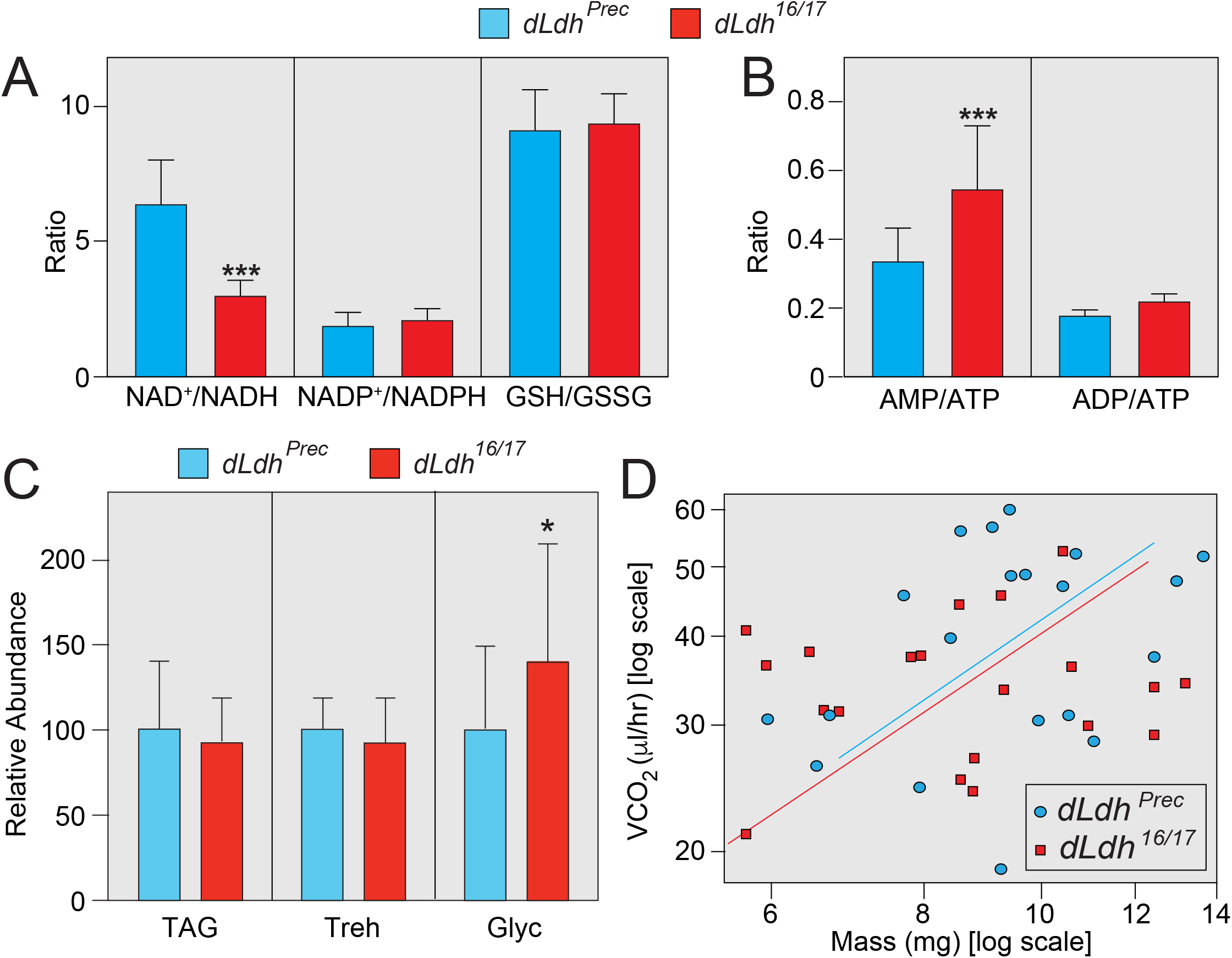
dLDH maintains the NAD^+^/NADH redox balance during larval development. Targeted LC-MS/MS analysis was used to measure metabolites associated with redox balance in *dLdh^prec^* controls and *dLdh^16/17^* mutants. The ratio of (A-B) NAD^+^/NADH, NADP^+^/NADPH, GSH/GSSG, AMP/ATP and ADP/ATP were determined in control and mutant larvae. n=8 samples collected from independent populations with 100 mid-L2 larvae per sample. (C) *dLdh^prec^* controls and *dLdh^16/17^* mutants were collected as mid-L2 larvae and the concentration of triglycerides (TAG), trehalose (Treh), and glycogen (Glyc) were measured in whole animal homogenates. All data were normalized to soluble protein. For all assays, n>10 samples collected from independent populations with 25 mid-L2 larvae per sample. (D) The rate of CO_2_ production was measured *dLdh^16/17^* mutants and precise excision controls. For (A-C), * *P* < 0.05. *** *P* < 0.001. Error bars represent one standard deviation.

Despite the fact that redox balance is significantly altered in *Ldh* mutants, the phenotypic consequences of this metabolic disruption are mild. We previously demonstrated that *dLdh* mutant larvae raised under ideal culture conditions can grow at a normal rate for much of larval development (Li et al., 2017). Furthermore, *dLdh* mutants that survive the mid-L3 lethal phase develop into adults (Li et al., 2017). To determine if any other biosynthetic processes are disrupted in *dLdh* mutants, we quantified the major larval pools of stored energy. Our analysis revealed that loss of dLDH activity has no effect on either triglyceride or trehalose levels (Figure 1C). Meanwhile, glycogen levels exhibited a modest, but significant increase in *dLdh* mutants when compared with control larvae (Figure 1C). This later observation was notable because the epidermis of human GSD-XI patients also appear to accumulate excess glycogen (Yoshikuni et al., 1986), indicating that *dLdh* mutants phenocopy the subtle metabolic defects observed in humans lacking LDHA.

### G3P levels are elevated in *dLdh* mutants

The ability of *dLdh* mutants grow at a normal rate suggests that *Drosophila* development adapts to the loss of dLDH activity. In this regard, human cell culture studies suggest that LDH-A inhibition can increase flux from glycolysis into the tricarboxylic acid cycle (TCA cycle) (Billiard et al., 2013; Xie et al., 2014). We found no evidence, however, that this metabolic shift occurs in flies, as *dLdh* mutant and control larvae produced CO_2_ at similar rates (Figure 1D). Furthermore, mitochondrial DNA content was unchanged in *Ldh* mutants (Supplemental Figure 1A), suggesting that loss of dLDH activity does not induce excess mitochondrial proliferation. Since our initial metabolic characterization failed to provide an adequate explanation for how larvae compensate for the loss of LDH, we turned to an untargeted metabolomics approach to poll a larger pool of analytes. Four independent GC-MS-based studies revealed that *dLdh^16/17^* mutants exhibit reproducible changes in only four metabolites: lactate, pyruvate, 2-hydroxyglutarate (2HG), and G3P (Figure 2A-B, Supplemental Tables 2-6). Since previous studies have already examined the relationship between dLDH and the metabolites lactate, pyruvate, and 2HG levels (Li et al., 2017), we focused our efforts on understanding why the G3P pool size was increased in *dLdh* mutants. To confirm that *dLdh* mutants accumulate excess G3P as the result of decreased dLDH activity, we demonstrated that expression of an *dLdh* transgene (*p{dLdh}*) in *dLdh* mutant larvae can restore G3P levels to those observed in *dLdh^prec^* control larvae (Figure 2C). Similarly, ubiquitous expression of a *UAS-dLdh-RNAi (dLdhi*) transgene induced elevated G3P levels (Figure 2D), thus confirming that *Drosophila* larvae accumulate excess G3P in response to the loss of dLDH activity. Considering that G3P levels are elevated in both GSD-XI patients and pancreatic cancer cells exposed to an LDH inhibitor ((Billiard et al., 2013; Boudreau et al., 2016; Miyajima et al., 1995)), our findings suggest that both flies and humans accumulate G3P to compensate for the loss of LDH activity.

**Figure 2.**
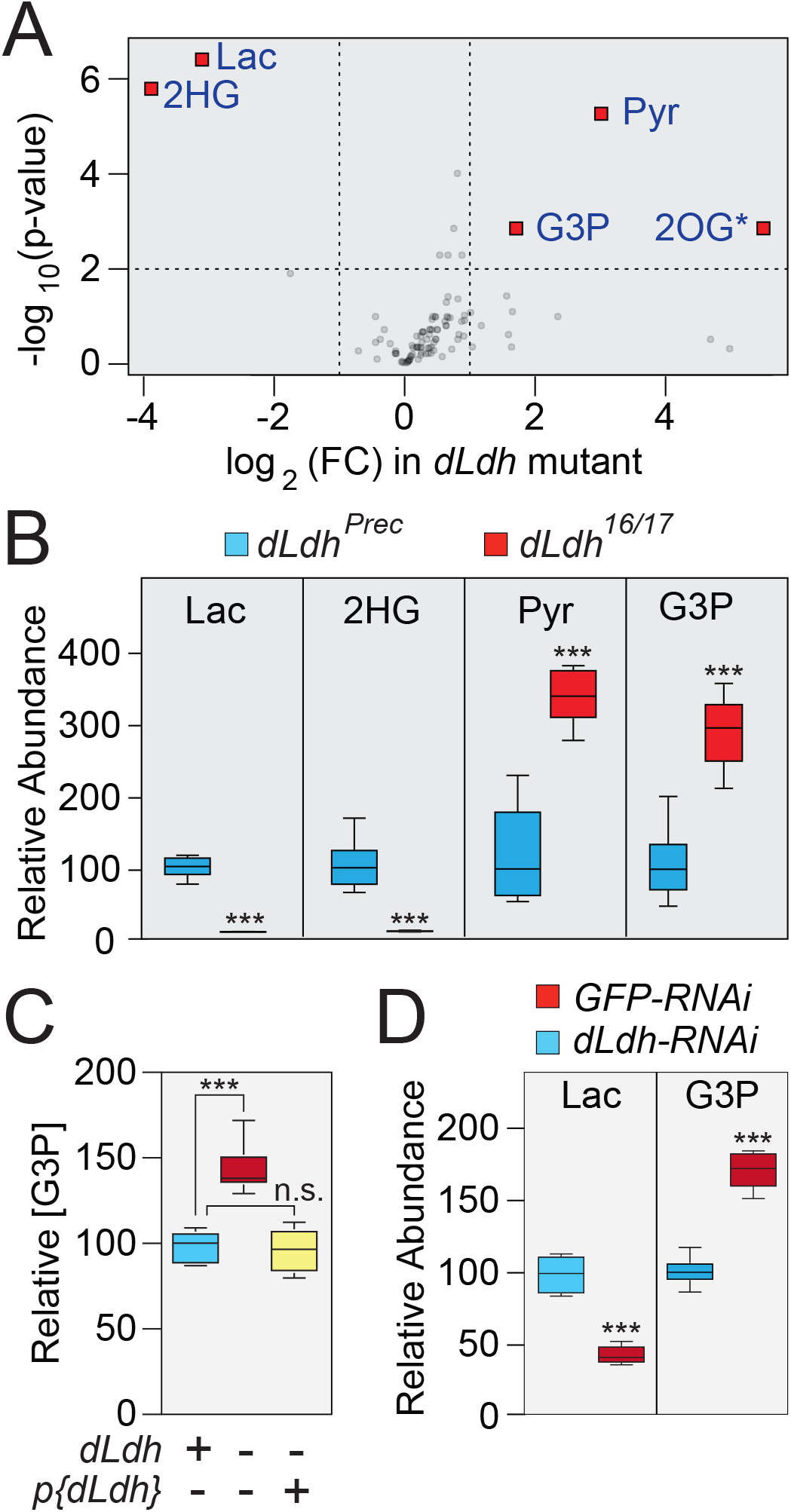
Metabolomic analysis of *dLdh* mutants. Data from GC-MS metabolomic analysis (Supplemental Table 3) comparing *dLdh^prec^* controls and *dLdh^16/17^* mutants were analyzed using Metaboanalyst. (A) A volcano plot highlighting metabolites that exhibited a >1.5-fold change and a p-value of <0.01. *Note that changes in 2OG levels were not reproducible is subsequent experiments. (B) The relative abundance of metabolites that exhibited significant changes in all four GC-MS experiments (*P* < 0.01; see Supplemental Table 2). (C) The relative abundance of G3P was measured in *dLdh^prec^* controls, *dLdh^16/17^* mutants, and *p{dLdh}; dLdh^16/17^* rescued animals during the L2 larval stage. (D) Lactate and G3P levels were measured in L2 larvae that ubiquitously expressed either a *UAS-GFP-RNAi* construct or a *UAS-dLdh-RNAi* construct under the control of *da-GAL4*. Abbreviations as follow: lactate (Lac), 2-hydroxyglutarate (2HG), pyruvate (Pyr), glycerol-3-phosphate (G3P), 2-oxoglutarate (2OG), not significant (n.s.). n=6 samples collected from independent populations with 25 mid-L2 larvae per sample. *** *P* < 0.001.

### GPDH1 regulates larval NAD^+^/NADH redox balance

Since GPDH1 couples G3P synthesis to NADH oxidation, increased G3P production could provide *dLdh* mutants with an alternative means of regenerating NAD^+^ (Figure 3A). Moreover, GPDH1 is a highly abundant protein in *Drosophila* larvae and could itself represent a key regulator of larval redox balance. To test these possibilities, we generated two *Gpdh1* loss-of-function alleles, *Gpdh1^A10^* and *Gpdh1^B18^*, both of which represent frameshift mutations that either delete or truncate the C-terminal catalytic domain, which is required for GPDH1 enzyme activity (Supplemental Figure 2A-B). Larvae that harbor a trans-heterozygous combination of these alleles, *Gpdh1^A10/B18^*, exhibited significantly lower G3P levels compared to controls (Figure 3B). Furthermore, ubiquitous expression of a *UAS-Gpdh1* transgene restored normal G3P levels in mutant larvae (Figure 3B), confirming that the loss of zygotic GPDH1 reduces G3P synthesis.

**Figure 3.**
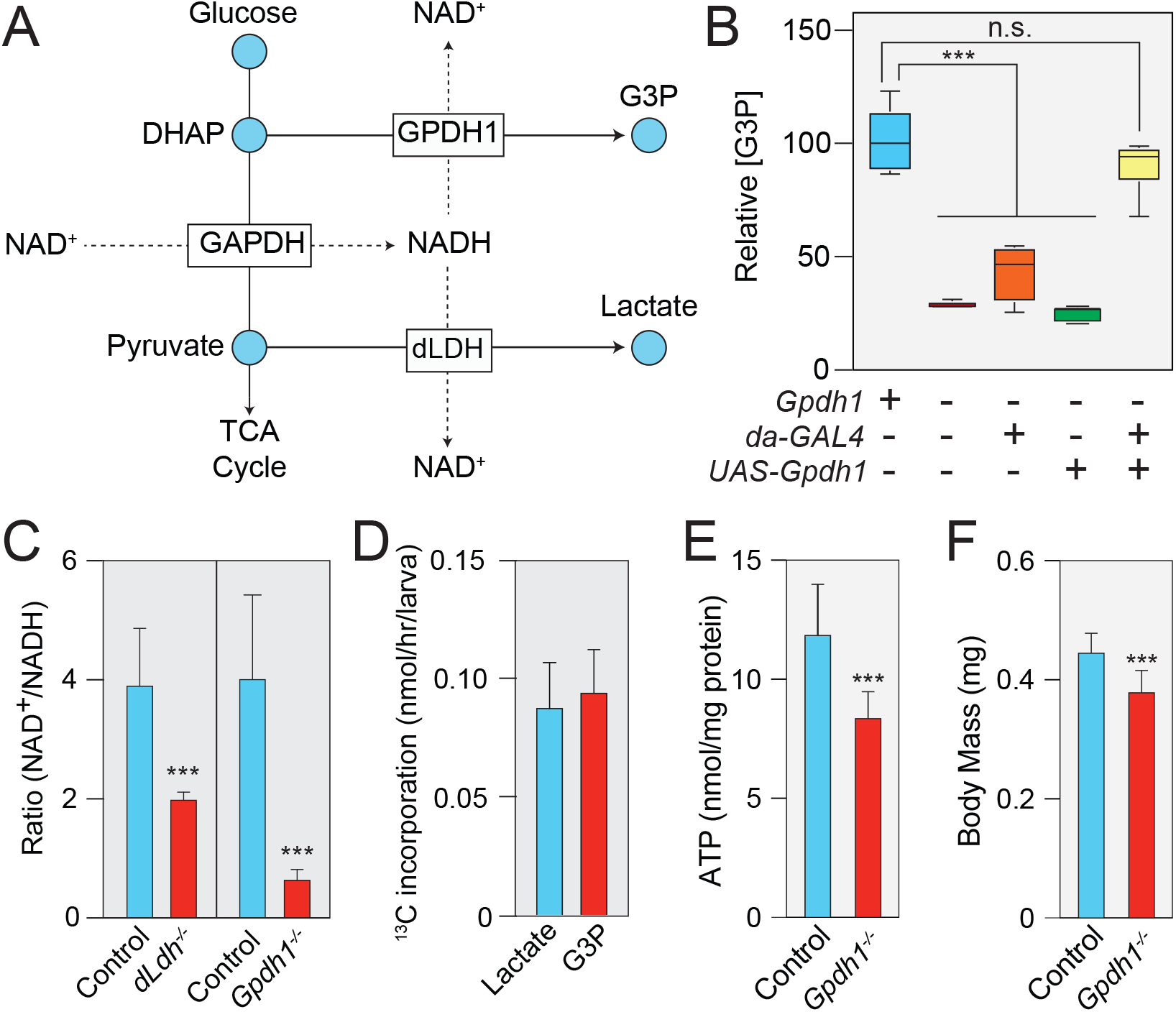
GPDH1 controls NAD^+^/NADH redox balance during larval development. (A) A schematic diagram illustrating how GPDH1 and dLDH redundantly influence NAD^+^ levels. Abbreviations as follow: dihydroxyacetone phosphate (DHAP), glycerol-3-phosphate (G3P), pyruvate (Pyr), lactate (Lac). (B) GC-MS was used to measure relative G3P abundance in mid-L2 larvae for the following five genotypes: *Gpdh1*^*A10/*+^, *Gpdh1^A10/B18^, Gpdh1^A10/B18^; da-GAL4, Gpdh^AW/B18^; UAS-Gpdh1*, and *Gpdh1^A10/B18^; da-GAL4 UAS-Gpdh1*. Data are represented as box plots, *n* = 6. \*\*\**P* < 0.001. (C) The NAD^+^/NADH ratio in mid-L2 larvae of the following genotypes: *dLdh^prec^*, *dLdh^16/17^*, *Gpdh1*^*A10/*+^, and *Gpdh1^A10/B18^*. (D) mid-L2 larvae were fed D-glucose-^13^C_6_ for two hours and the rate of ^13^C isotope incorporation into lactate (Lac) and G3P was determined based on m+3 isotopologue abundance. (E) ATP levels are significantly decreased in *Gpdh1^A10/B18^* as compared with *Gpdh1*^*A10/*+^ controls. (F) The body mass of *Gpdh1^A10/B18^* larvae is significantly lower than that of *Gpdh1*^*A10/*+^ controls 0-4 hours after the L2-L3 molt. In panels (B-F), n=6 samples per genotype. *\*\*\*P* < 0.001. For (C-F), error bars represent one standard deviation.

To determine if G3P production influences larval redox balance, we measured both NAD^+^ and NADH levels in *Gpdh1* mutants. Similar to *dLdh* mutants, we observed that the NAD^+^/NADH ratio in mid-L2 larvae was significantly lower in *Gpdh1* mutants as compared with a *Gpdh1*^*A10/*+^ heterozygous control (Figure 3C). We also examined the extent to which GPDH1 influences NAD^+^ levels by feeding D-glucose-^13^C_6_ to mid-L2 larvae and measuring the rate of lactate and G3P synthesis. Since both dLDH and GPDH1 must oxidize one molecule of NADH in order to form one molecule of either lactate or G3P respectively (Figure 3A), we can indirectly infer the rate at which each enzyme regenerates NAD^+^ based on synthesis of these metabolites. Our analysis revealed that mid-L2 larvae synthesize m+3 lactate and m+3 G3P at similar rates (Figure 3D), indicating that GPDH1 and dLDH regenerate roughly equivalent amounts of NAD^+^ during larval development.

Our observation that *Gpdh1* mutants exhibit a decreased NAD^+^/NADH ratio balance also implicates this enzyme in coordinating redox balance with larval energy production. G3P synthesized in the cytoplasm can be oxidized on the inner mitochondrial membrane by the FAD dependent enzyme Glycerophosphate oxidase 1 (Gpo-1; FBgn0022160) to generate ATP via oxidative phosphorylation (O’Brien and MacIntyre, 1972). Therefore, any decrease in G3P synthesis could also reduce ATP production. Consistent with this function, ATP levels were significantly decreased in *Gpdh1* mutants when compared with controls (Figure 3E). Yet, despite the role for GPDH1 in these critical metabolic processes, animals lacking zygotic GPDH1 activity exhibit only mild developmental defects. In agreement with previous reports (Bewley and Lucchesi, 1977), we found that *Gpdh1* mutants are viable through larval development and are ~15% smaller at the L2-L3 molt (Figure 3F and Supplemental Figure 2C), a result which again demonstrates how larval development is robust and can compensate for significant metabolic insults.

### *Gpdh1; dLdh* double mutants exhibit severe developmental delays and a synthetic lethal phenotype

Since dLDH and GPDH1 individually control larval redox balance, we tested the possibility that simultaneous removal of both enzymes would induce growth defects. Indeed, when compared with *Gpdh1* and *dLdh* single mutants, *Gpdh1; dLdh* double mutants are 85% smaller and experience developmental delays (Figure 4A-C). Moreover, *Gpdh1; dLdh* double mutants die throughout L1 and L2 development, with ~30% of double mutant larvae dying during L1 development and all animals failing to complete the L2-L3 molt (n>100; Figure 4C). To further characterize these growth defects in double mutant larvae, we examined the larval brain, which was previously reported to display high levels of both dLDH and GPDH1 activity (Rechsteiner, 1970). *w^1118^* controls, *Gpdh1* and *dLdh* single mutants, and *Gpdh1; dLdh* double mutants were collected 60 hours after egg-laying and the brains were fixed and stained with DAPI, to visualize overall tissue size, and for Deadpan (Dpn), to visualize neuroblasts (Figure 4D-H). While the brains of single mutant larvae exhibited no growth defects, the brains of *Gpdh1; dLdh* double mutants were significantly smaller than controls (Figure 4D-G). However, if *Gpdh1; dLdh* double mutants were allowed to develop for an additional 24 hours, the brain grew to a comparable size as the *w^1118^* control (Figure 4D,H). We observed a similar phenomenon in the larval intestine, where the posterior midgut of age-matched *Gpdh1; dLdh* mutant larvae was shorter than either single mutant, contained significantly smaller enterocyte nuclei, and exhibited a growth delay of approximately 24 hours when compared with single mutant controls (Supplemental Figure 3). To determine if these growth phenotypes are associated with a decreased rate of cell cycle progression, we dissected brains from size-matched larvae and incubated them in an organ culture media with EdU for 2 hours. When compared with *w^1118^* controls, *dLdh* single mutants, and *Gpdh1* single mutants, the number of cells that stain with EdU was significantly decreased in *Gpdh1; dLdh* mutant brains (Figure 5A-E). Overall, these results demonstrate that while larval development can compensate for the loss of either dLDH or GPDH1, removal of both enzymes severely restricts tissue growth.

**Figure 4.**
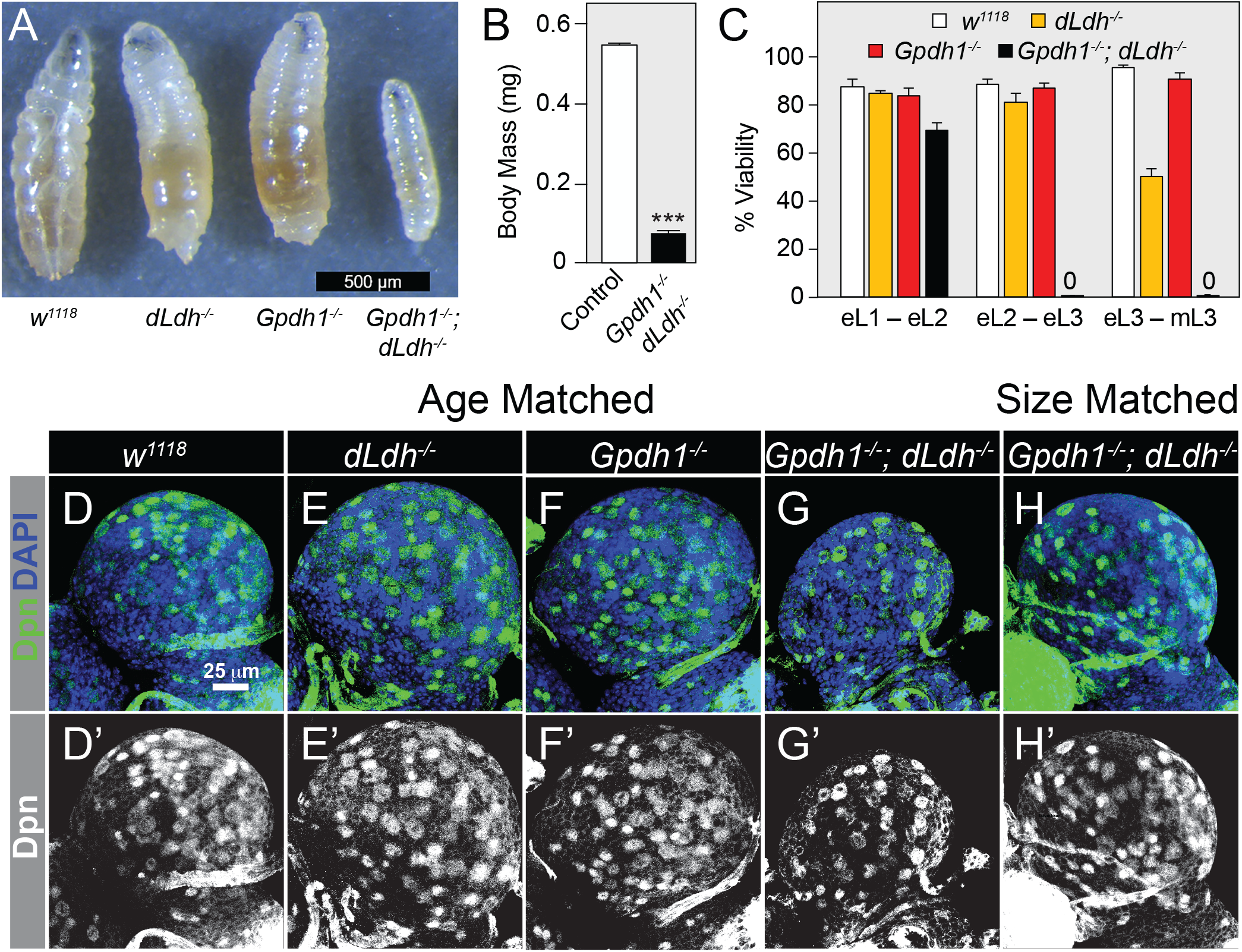
*Gpdh1; dLdh* double mutants exhibit severe growth phenotypes. (A) Representative images of larvae from synchronized populations of w^1118^, *dLdh^16/17^, Gpdh1^A10/B18^*, or *Gpdh1^A10/B18^; dLdh^16/17^* double mutants. Note the severe developmental delay exhibited by double mutant larvae. (B) The body mass of *w^1118^* and *Gpdh1^A10/B18^; dLdh^16/17^* double mutant larvae measured 72 hours after egg-laying. (C) The viability of the four genotypes listed in (A) were measured during defined periods of larval development. (B,C) Error bars represent one standard deviation. n=6 samples per genotype. (D-H) Maximum projections of dorsal half of L2 larval brains stained for Dpn (green) and DAPI (blue) from (D) *w^1118^* controls, (E) *dLdh^16/17^* mutants, (F) *Gpdh1^A10/B18^* mutants, (G) age-matched *Gpdh1^A10/B18^; dLdh^16/17^* double mutants, and (H) size-matched *Gpdh1^A10/B18^; dLdh^16/17^* double mutants. The scale bar in (D) applies to (E-H). Note that the (D’-H’) display the Dpn channel alone in gray scale. \*\*\**P* < 0.001.

**Figure 5.**
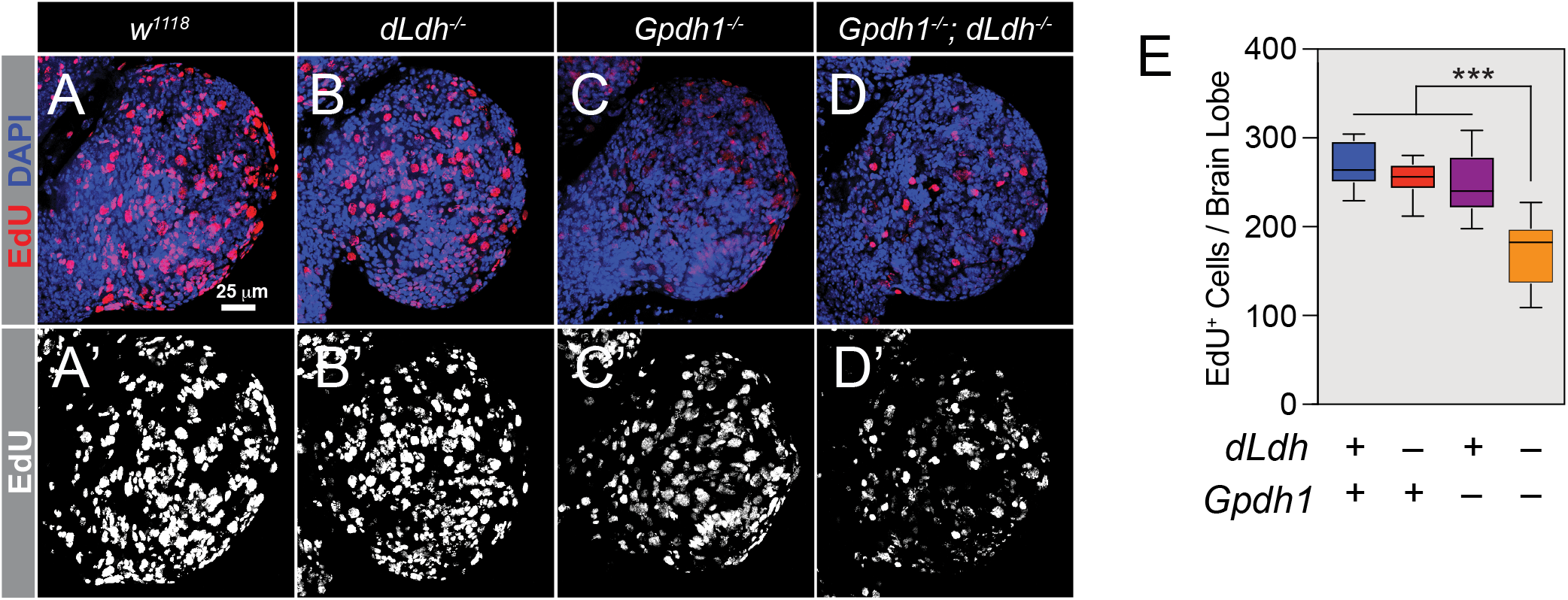
The DNA replication rate is decreased in brains of *Gpdh1; dLdh* double mutants. (A-D) Maximum projections of dorsal half of size-matched L2 larval brains stained for EdU (red) and DAPI (blue) from (A) *w^1118^* controls, (B) *dLdh^16/17^* mutants, (C) *Gpdh1^A10/B18^* mutants, and (D) *Gpdh1^A10/B18^; dLdh^16/17^* double mutants. The scale bar in (A) applies to (A-D). Panels (A’-D’) display EdU staining alone in gray scale. (E) Histogram of the number of EdU positive cells per dorsal brain lobe per genotype. For all panels, p-value adjusted for multiple comparisons using the Bonferroni-Dunn method. * *P* < 0.05. **P<0.01, \*\*\**P* < 0.001.

### Simultaneous loss of dLDH and GPDH1 induces defects in carbohydrate and amino acid metabolism

Since our metabolic studies suggested that dLDH and GPDH1 maintain the larval NAD^+^/NADH ratio, we next examined the possibility that the *Gpdh1; dLdh* double mutant growth phenotypes stem from a severe disruption of redox balance. These experiments revealed, however, that the ratio of NAD^+^ to NADH was similar in control and double mutant larvae (Figure 6A). To further investigate this unexpected result, we used a targeted GC-MS-based approach to analyze central carbon metabolism of both *Gpdh1* single mutants and *Gpdh1; dLdh* double mutants (Figure 6B-D; Supplemental Tables 7 and 8). In the case of the *Gpdh1* mutant larvae, metabolomic analysis revealed a significant disruption of amino acid metabolism. Not only were aspartate and several essential amino acids decreased, but we also observed elevated levels of urea and the urea cycle intermediate ornithine (Figure 5B-D), suggesting that loss of GPDH1 results in elevated amino acid catabolism. Moreover, our analysis also uncovered elevated glutamate and proline levels. Considering that insects can synthesize proline in a NADH dependent manner (Mccabe and Bursell, 1975), this finding hints at a mechanism by which loss of G3P production induces elevated proline synthesis in response to aberrant redox balance. Intriguingly, *Gpdh1* mutants do not exhibit an increase in either lactate or 2HG levels (Figure 6B,C) – a result which would support the inverse correlation between GPDH1 and dLDH activity. This null result, however, could indicate that dLDH activity is saturated in developing larvae. Finally, xanthine and urate levels, which are produced by purine catabolism, were also increased in *Gpdh1* mutants (Figure 6B; Supplemental Figure 4A).

**Figure 6.**
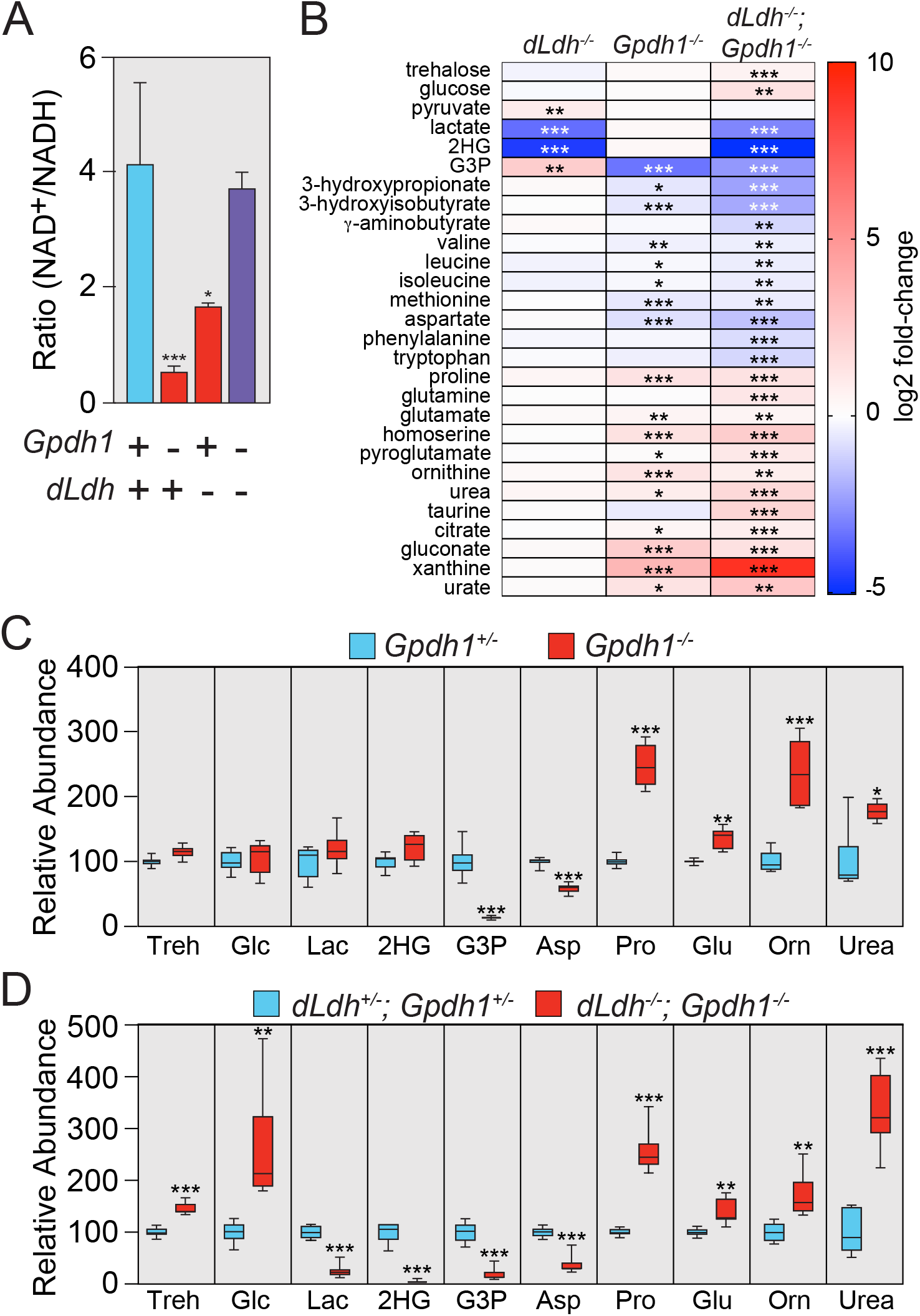
Amino acid and glucose metabolism are disrupted in *Gpdh1; dLdh* double mutants. (A) The NAD^+^/NADH ratio was measured in *w^1118^, dLdh^16/17^, Gpdh1^A10/B18^*, and *Gpdh1^A10/B18^; dLdh^16/17^* mid-L2 larvae. Error bars represent one standard deviation. n=6 samples per genotype. (B) A heat-map summarizing changes in metabolite abundance in *dLdh^16/17^* mutants relative to *dLdh^prec^* controls, *Gpdh1^A10/B18^* mutants relative to *Gpdh1*^*A10*/+^ controls, and *Gpdh1^A10/B18^; dLdh^16/17^* double mutants relative to *Gpdh1*^*A10/*+^*; dLdh^16/+^* controls. The abundance of select metabolites for either (C) *Gpdh1^A10/B18^* mutants relative to *Gpdh1*^*A10*/+^ controls or (D) *Gpdh1^A10/B18^; dLdh^16/17^* double mutants relative to *Gpdh1^A10/+^; dLdh^16/+^* are represented as box plots. For all panels, p-value adjusted for multiple comparisons using the Bonferroni-Dunn method. * *P* < 0.05. **P<0.01, \*\*\**P* < 0.001.

Nearly all of the metabolic changes observed in the *Gpdh1* single mutant were enhanced in the *Gpdh1; dLdh* double mutants (Figure 6B,D). For example, *Gpdh1; dLdh* mutant larvae exhibited a 500-fold increase in xanthine levels when compare to the heterozygous controls (Figure 6B; Supplemental Figure 4B), suggesting a severe disruption of purine metabolism. Overall, the metabolic changes observed in the double mutant represented the combined metabolic disruptions seen in the single mutants (Figure 6B-D), but with two major exceptions – the relative abundance of both trehalose and glucose were significantly elevated in *Gpdh1; dLdh* mutant larvae (Figure 6B,D). This result is important as it suggests that the loss of both enzymes lead to decreased glycolytic flux. Since an inhibition of glucose catabolism would result in decreased NADH formation, the *Gpdh1; dLdh* mutant metabolomic profile provides an explanation for why the NAD^+^/NADH ratio is normal in double mutants. Moreover, we also observed that ATP levels are dramatically decreased in *Gpdh^A10/B18^; dLdh^16/17^* double mutants when compared with *Gpdh^A10/+^; dLdh^16/+^* heterozygous controls (Supplemental Figure 4C). This result demonstrates that loss of both enzymes limits ATP production and is consistent with a model in which glycolytic flux is restricted in double mutant larvae. Overall, our metabolomic approach not only demonstrates that the growth defects caused by loss of both dLDH and GPDH1 are associated with severe disruption of central carbon metabolism but also highlights the plasticity of animal metabolism.

## DISCUSSION

Our findings demonstrate that the ability of dLDH and GPDH1 to cooperatively regulate NAD^+^/NADH redox balance and carbohydrate metabolism imparts robustness on larval growth. This relationship likely serves multiple purposes, as the production of lactate and G3P metabolism not only influences larval NAD^+^/NADH redox balance, but also controls the pool size of glycolytic intermediates and dictates the manner by which cells generate ATP. Considering that human cells also up-regulate GPDH1 activity in response to decreased lactate synthesis, our findings indicate that this metabolic relationship is conserved across animal phyla and hint at a mechanism by which GPDH1 activity could render tumors resistant to LDH inhibitors.

### The roles of LDH and GPDH1 in cancer and animal development

The possibility of using LDH inhibitors to disrupt tumor growth was first proposed over 60 years ago, shortly after the discovery that the pyruvate analog oxamate disrupts aerobic glycolysis and slows the growth of HeLa cells (Goldberg and Colowick, 1965; Papaconstantinou and Colowick, 1961). During the last decade, the goal of using LDH inhibitors as chemotherapeutic agents has been revisited, with several studies demonstrating that this approach can disrupt cancer cell growth (Billiard et al., 2013; Boudreau et al., 2016; Daniele et al., 2015; Fantin et al., 2006; Qing et al., 2010). Yet, despite the promise of such compounds, studies of human and mouse *LDHA* mutants raise concerns about the potential effectiveness of inhibiting LDH. First, GSD-XI patients grow and develop normally (Kanno et al., 1988; Kanno et al., 1980; Miyajima et al., 1995), suggesting that human developmental metabolism can compensate for loss of this enzyme. Secondly, although *LDHA* inhibition induces elevated TCA cycle flux in cell culture, this reliance on the TCA cycle is not observed in neither tumors derived from conditional *LDHA* mutant nor *ex vivo* tumor slices treated with an LDH inhibitor (Xie et al., 2014). Such observations are important because they suggest that the metabolic plasticity of cells in culture differs significantly from tissues *in vivo*.

Our studies in the fly support the *in vivo* mammalian observations – *dLdh* mutants grow normally and do not increase CO_2_ production, indicating that flux through the TCA cycle is unchanged. Instead, we observed that *dLdh* mutants specifically up-regulate G3P synthesis as a means of maintaining developmental growth. This finding is consistent with decades of observation in tumors, insects, and healthy human tissues, which, on the whole, repeatedly pointed to an inverse correlation between lactate and G3P production (Boxer and Shonk, 1960; Miyajima et al., 1995; Rechsteiner, 1970; Zebe and McShan, 1957). Moreover, recent cell culture studies have also demonstrated that LDH inhibitors induce G3P synthesis, thus demonstrating that this metabolic relationship is present in cultured cells (see supplemental data in Billiard et al., 2013; Boudreau et al., 2016). Overall, our observations in the fly suggest a common metabolic relationship that allows animal cells to adapt to redox stress.

The link between larval redox balance and the role of G3P in ATP production could also explain a contradiction in the *Drosophila* metabolism literature. Mutations that disrupt either glycolysis (e.g. *dERR, Pfk*) or the electron transport chain (ETC) result in severe growth defects (Mandal et al., 2005; Meiklejohn et al., 2013; Tennessen et al., 2011). In contrast, larvae that harbor mutations in either the *mitochondrial pyruvate carrier (dMPC1*) or *malate dehydrogenase 2* are able to complete larval development with relatively mild phenotypes (Bricker et al., 2012; Wang et al., 2010). Similarly, larvae that lack zygotic *isocitrate dehydrogenase 3b* exhibit developmental delays but are able to survive until metamorphosis (Duncan et al., 2017). These observations suggest that while glycolysis and oxidative phosphorylation are necessary for development, larvae do not require a fully functional TCA cycle. This arrangement makes sense in that larval metabolism is largely dedicated to shuttling metabolic intermediates into biosynthetic pathways. By activating GPDH1, the production of G3P helps regenerate cytosolic NAD^+^ without increasing CO_2_ production while also allowing cells to transfer reducing equivalents to the ETC and generate ATP.

### *Drosophila* as a model for studying metabolic plasticity

Our study highlights the remarkable metabolic plasticity that underlies animal development and physiology. Intermediary metabolism adapts to a surprisingly broad range of natural genetic variation, dietary stress, and metabolic insults. For example, mutations in the *Drosophila* mitochondrial pyruvate carrier *dMPC1*, which render cells unable to transport pyruvate into the mitochondria, elicit no obvious phenotypes when mutant larvae are raised under standard growth conditions (Bricker et al., 2012). Moreover, natural populations of *Drosophila* can buffer larval development against significant variations in mitochondrial oxidative capacity and the scaling relationship between mass and metabolic rate (Matoo et al., 2018). A similar phenomenon is also observed in *C. elegans*, where entire metabolic pathways are rewired in response to dietary stress or genetic mutations (MacNeil et al., 2013; Watson et al., 2014; Watson et al., 2016). These examples not only highlight the adaptability of animal metabolism, but also emphasize how little we understand about this topic. The molecular mechanisms that control adaptive metabolic rewiring, however, are often difficult to study in a laboratory setting, where animals are raised on high nutrient diets and buffered against the environmental stress. In this regard, recent advances in metabolomics provides a powerful approach for understanding how metabolism adapts to environmental and genetic insults. By analyzing changes in gene expression within the context of metabolomic data, compensatory changes in metabolic flux can be quickly identified and analyzed using standard model organism genetics. The power of this approach is demonstrated by our studies of *dLdh*. Even without prior knowledge of the link between LDH and GPDH1 activity, our study pinpointed increased G3P synthesis as the adaptive response within *dLdh* mutants, thus demonstrating how metabolomics holds the potential to illuminate the complex metabolic network that supports animal development.

## ACKNOWLEDGEMENTS

We thank the Bloomington *Drosophila* Stock Center (NIH P40OD018537) for providing fly stocks, the *Drosophila* Genomics Resource Center (NIH 2P40OD010949) for genomic reagents, and Flybase (NIH 5U41HG000739). Targeted GC-MS analysis was conducted using instruments housed in the Indiana University Mass Spectrometry Facility, which is supported, in part, by NSF MRI Award 1726633. Some of the metabolomic analyses described herein performed at the Metabolomics Core Facility at the University of Utah, which is supported by 1S10OD016232-01, 1S10OD021505-01 and 1U54DK110858-01. Special thanks to Madhulika Rai for useful comments. C.J.G. was supported, in part, by the Lawrence University LU-R1 program. K.B. and N.S.S. were supported by National Institute of Health Award R01GM124220. C.R.J. was supported by NSF CAREER award to K.L.M. (IOS-1149178, IOS-1505247). J.M.T. is supported by the National Institute of General Medical Sciences of the National Institutes of Health under a R35 Maximizing Investigators’ Research Award (MIRA; 1R35GM119557).

## Supplemental Figure Legends

**Supplemental Figure 1.**
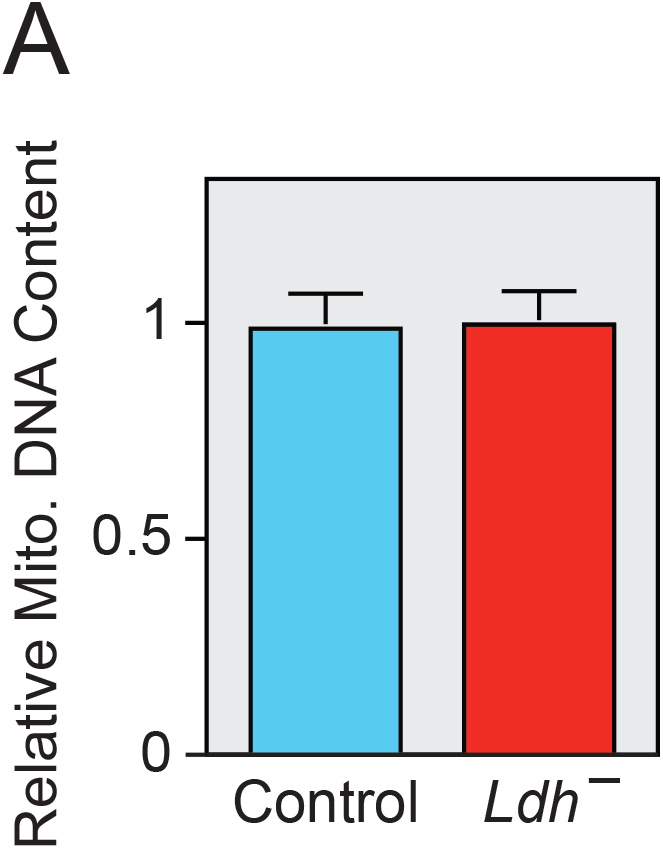
Mitochondrial abundance is unaffected by *dLdh* mutations. (A) The relative abundance of mitochondrial DNA is similar in *dLdh^prec^* controls and *dLdh^16/17^* mutants. Ratio is based on the abundance of *mt::CoI* copy number relative to *Rpl32* copy number. Error bars represent standard deviation.

**Supplemental Figure 2.**
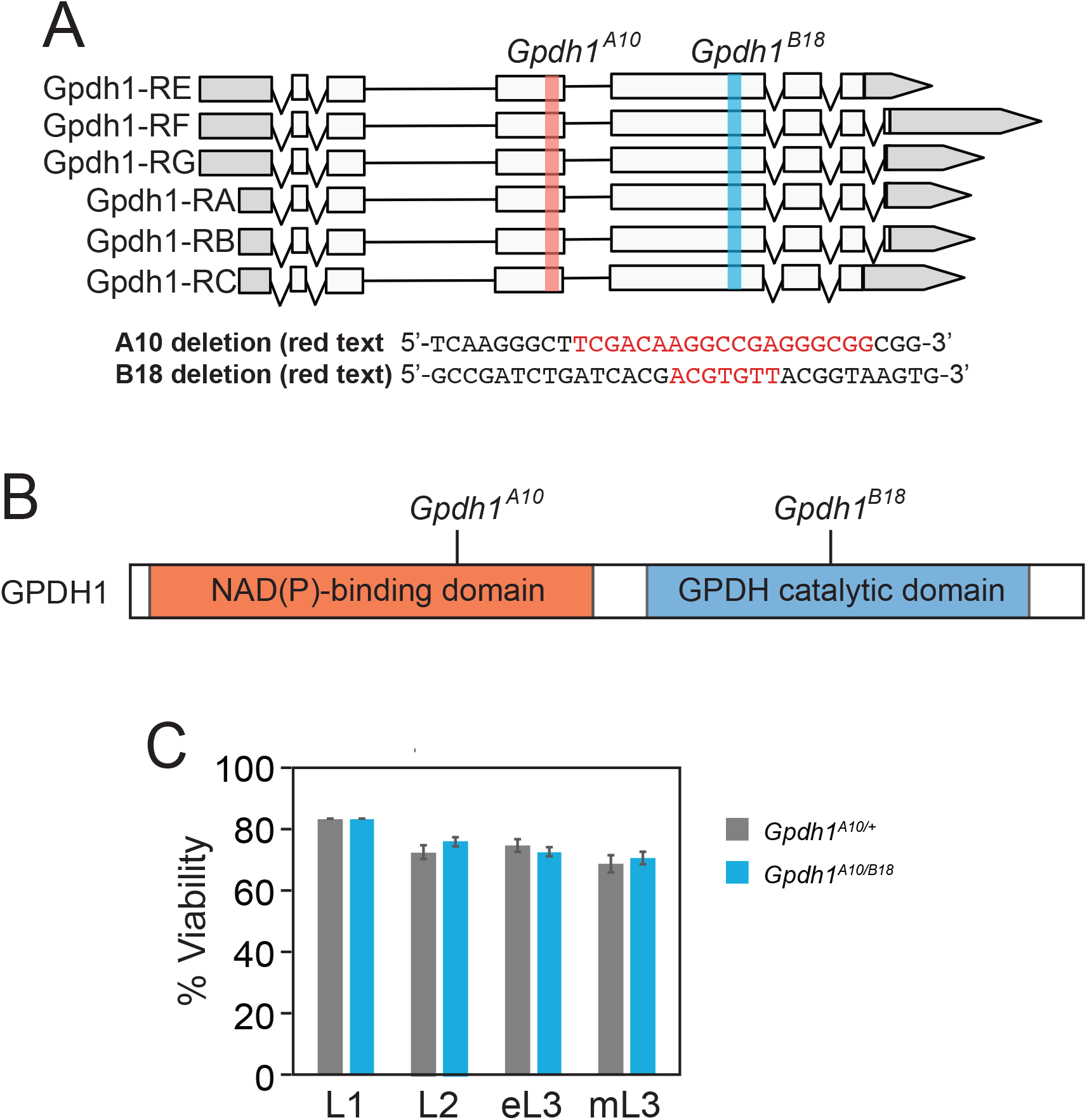
Generation of *Gpdh1* mutants. (A) A schematic diagram illustrating the *Gpdh1* locus, sequences targeted by guide RNA constructs, and sequence deleted by the *Gpdh1^A10^* and *Gpdh1^B18^* mutations. Deleted bases are highlighted in red. (B) A schematic diagram illustrating the location of the *Gpdh1^A10^* and *Gpdh1^B18^* mutations within the GPDH1 protein. (C) *Gpdh1*^*A10/*+^, and *Gpdh1^A10^/^B18^* mutants were analyzed for larval viability. Bars represent the percent of animals that survived from the previous developmental stage until the stage noted on the x-axis. Error bars represent standard deviation. n>100 larvae per timepoint.

**Supplemental Figure 3.**
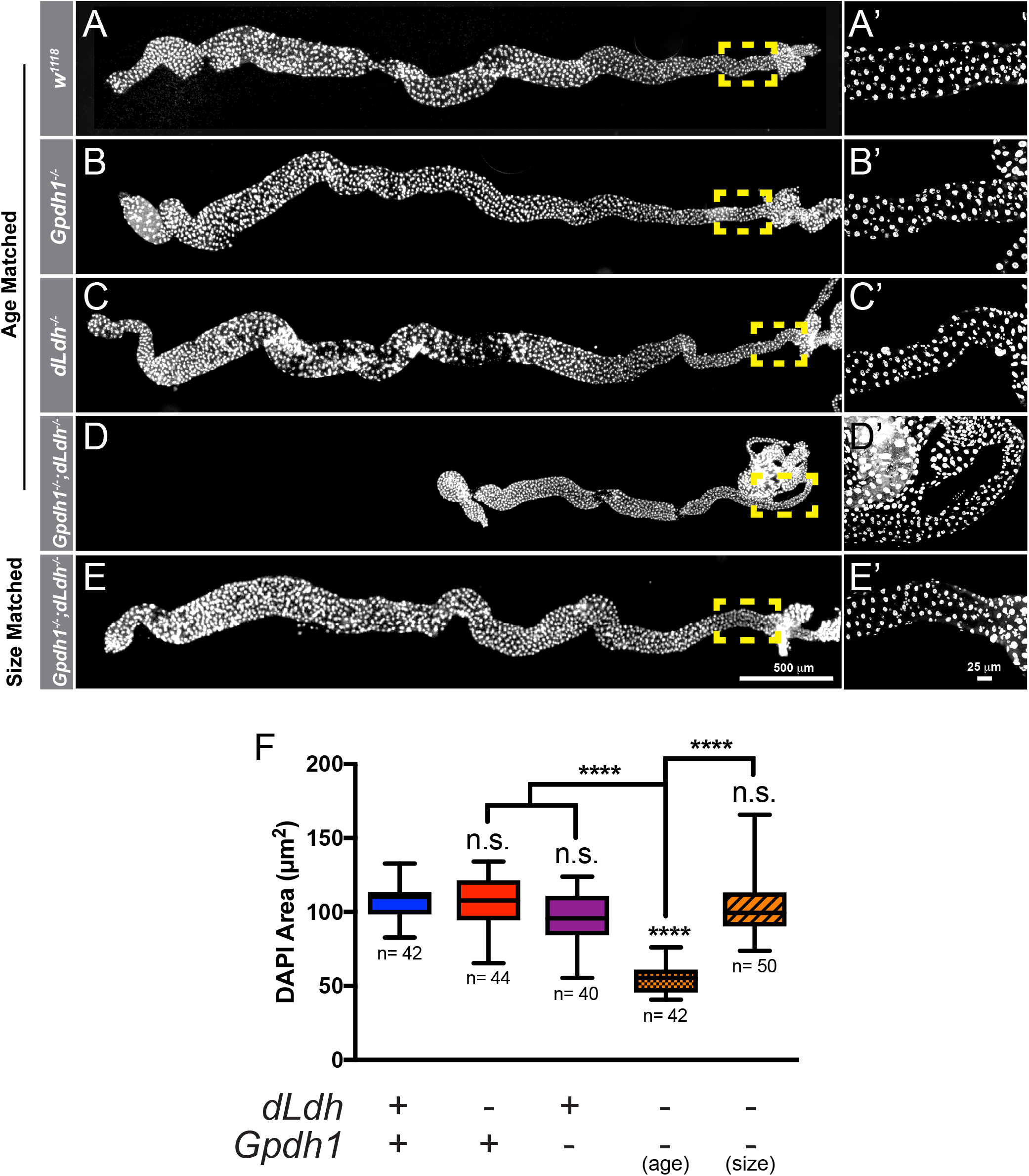
The intestine of *Gpdh^A10^/^B18^; dLdh^16/17^* double mutant larvae exhibit growth defects and contain smaller cells. (A-E) L2 larval posterior midguts (PMGs) stained for DAPI (gray) from (A) *w^1118^* controls, (B) *Gpdh1^A10/B18^* mutants, (C) *dLdh^16/17^* mutants, (D) age-matched *Gpdh1^A10/B18^; dLdh^16/17^* double mutants, and (E) size-matched *Gpdh1^A10/B18^; dLdh^16/17^* double mutants. (A’-E’) Magnified images of the outlined regions in A-E. The scale bar in (E) and (E’) apply to (A-E) and (A’-E’), respectively. (F) Histogram of nuclei size. **** P < 0.0001.

**Supplemental Figure 4.**
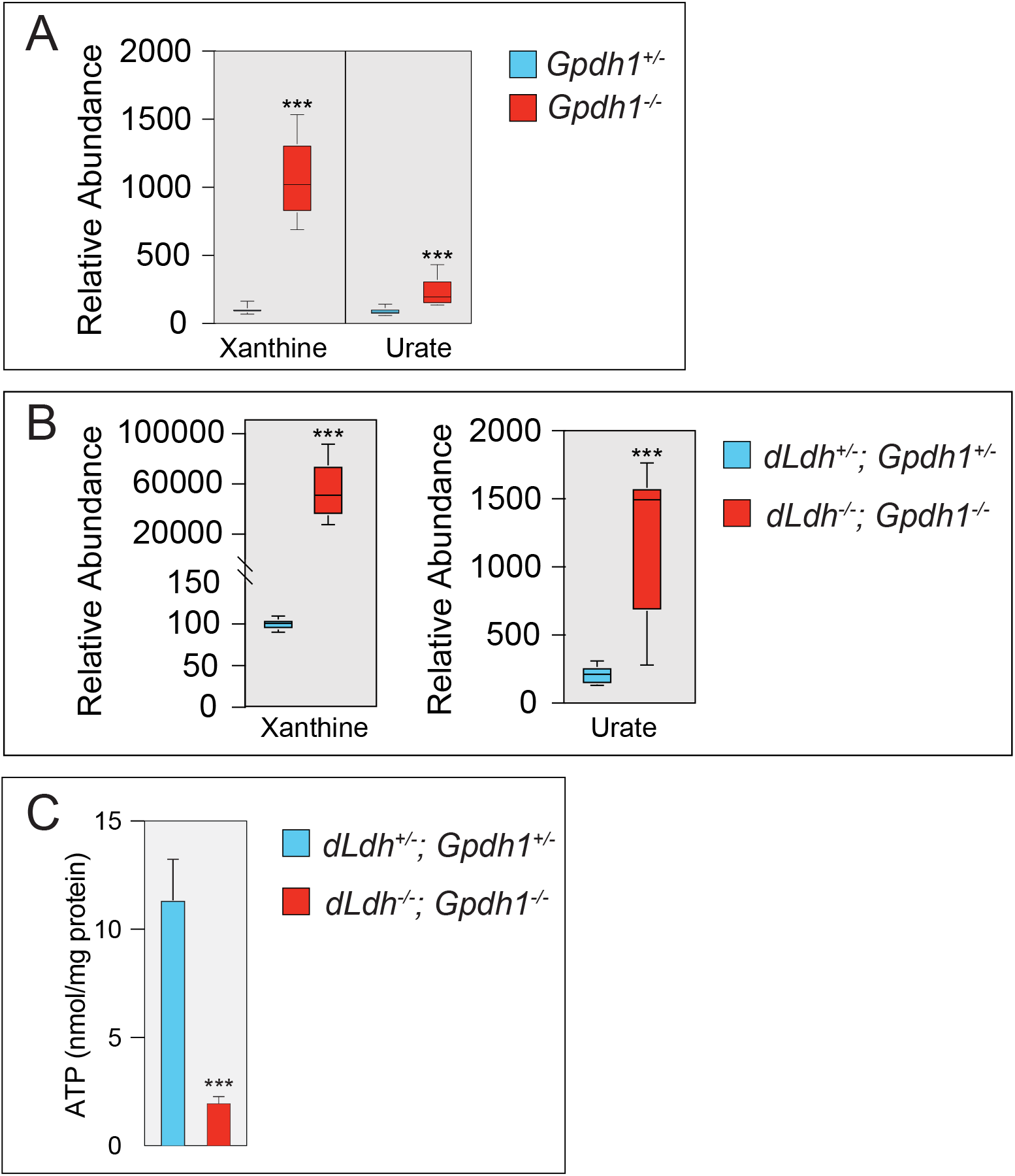
*Gpdh1; dLdh* double mutants exhibit significant changes in purine catabolism and ATP production. GC-MS was used to measure the relative abundance of xanthine and urate in either (A) *Gpdh1^A10^/^B18^* mutants relative to *Gpdh1^A10^/^+^* controls or (B) *Gpdh1^A10^/^B18^; dLdh^16/17^* double mutants relative to *Gpdh1^A10/+^; dLdh^16/+^*. (C) ATP levels are significantly decreased in *Gpdh1^A10^/^B18^; dLdh^16/17^* double mutants relative to *Gpdh1^A10/+^; dLdh*^16/+^. \*\*\**P* < 0.001.

